# Climate influences bill shape diversification in cavity-excavating birds

**DOI:** 10.1101/2022.09.20.508664

**Authors:** Vaibhav Chhaya, Sushma Reddy, Anand Krishnan

**Affiliations:** Department of Biology, Indian Institute of Science Education and Research (IISER) Pune, Pashan Road, Pune 411008, India; Bell Museum of Natural History and Department of Fisheries, Wildlife and Conservation Biology, University of Minnesota, St. Paul, MN, USA; Department of Biological Sciences, Indian Institute of Science Education and Research (IISER) Bhopal, Bhauri 462066, Madhya Pradesh, India

**Keywords:** computed tomography, geometric morphometrics, bill shape, barbets, morphological diversification

## Abstract

The remarkable structural and functional diversity of the avian bill serves as an ideal system to understand how ecological and environmental factors influence morphological diversification under physically demanding tasks. For example, in cavity-excavating birds where the bill experiences tremendous physical stresses, climate and habitat affect the mechanical properties of available nesting substrates and therefore can potentially influence the shape and performance of bills. Few studies have focused on the entire avian bill, a complex structure consisting of the rhamphotheca and bony core of the upper and lower mandibles, which may evolve as independent modules or as a single integrated unit. Here, we examined patterns and drivers of bill shape diversification in barbets using micro-computed tomography of museum study skins. We employed landmark-based geometric morphometrics and phylogenetic comparative methods to study the influence of climatic conditions, integration and phylogenetic constraints on bill shape diversification. Asian and African barbets have explored distinct maxillary shape spaces without corresponding change in mandibular shape after correcting for phylogenetic non-independence. Additionally, maxillary shape for both rhamphotheca and bony components was significantly correlated with climate variables and exhibited signatures of allometric scaling. All four bill components, however, covaried with each other and have evolved in a gradual and integrated fashion, exhibiting late-stage shape disparity. Through these findings, our study offers crucial insights into how ecological behaviors impact morphological evolution of complex, multifunctional features.

## INTRODUCTION

Morphological traits influence diet, movement and other ways that an organism interacts with its physical environment. By examining patterns of morphological trait diversification, we may understand both the evolutionary history of species and lineages, and the role that environmental changes play in shaping them. For example, organisms encountering ecological opportunity, such as those colonizing islands, undergo a rapid expansion in morphological diversity, accompanied by the occupation of new ecological and foraging niches (1–6). Alternatively, many speciose lineages exhibit morphological diversity that accumulates more gradually (7, 8). As a result of selective pressures acting either in concert with or independently of neutral processes, different morphological traits of a lineage diversify either independently or in correlation with each other (9). The concept of morphological integration suggests that certain morphological traits co-vary in evolutionary time owing to developmental or genetic constraints. Modularity, on the other hand, suggests that each module diversifies independently of others, and that the life-history of an organism emerges from the sum of these independently diversifying modules (10, 11). Studies have found evidence for both modularity and integration in diverse taxa, and sometimes support for a mix of both processes (12, 13, 22, 14–21). Much, however, remains to be understood about the role of morphological integration in driving form and function in complex features that serve diverse purposes.

The avian bill is a multifunctional appendage that serves as an excellent study in morphological and functional diversification. The bill of any bird consists of the upper maxilla and the mandible, each composed of a keratinous outer sheath (the rhamphotheca) and the inner bony structures (23–26). The evolution of this bill structure in birds, and the concomitant loss of teeth has led to a great expansion in both morphological diversity and function (27). Bird bills serve diverse functions such as manipulating food and substrates, performing courtship and territorial displays, building nests and regulating temperature (23, 28–32). Conventional theories hold that diet exerts a significant influence on bill shape in many taxa (1, 2, 33). However, multiple non-dietary factors may also influence bill functional traits (12), including biomechanical factors related to how the bill performs under various environmental conditions. Understanding the ecological and environmental conditions that bill shape has evolved in (for example, its relationship with climate, as a proxy for changes in ecology and the physical environment that tasks are performed in) provides important insights into the relationship between form and function. Additionally, bill shape may also diverge under phylogenetic and developmental constraints, for example, as a result of allometric scaling owing to changes in body size or developmental integration with the skull (12–14, 34).

Given the complexities of the form and function of the avian bill, there may be differences between the shape of rhamphotheca and the bony core of the bill, as well as variation between upper and lower jaw morphologies. Most studies of bill shape diversity have, however, treated the bill as a single unit of shape, and have not tested whether and how these separate components may be influenced by each other as well as subject to potentially differing ecological and evolutionary constraints. Further, these studies focus on the upper bill (maxilla) and typically consider only the shape of the outer rhamphotheca. Micro-computed tomography (μCT) of museum specimens now enables us to virtually dissect keratinous and bony bill structures (35), and we may thus examine the following questions: a) Are the maxilla and mandible of the bill modular or integrated in how they diversify? b) Are keratinous and bony bill structures integrated? c) Does bill shape diversification in birds exhibit a relationship to the external environment? and d) How do the different bill modules, allometric scaling and phylogenetic constraints influence bill shape in these lineages?

Here, we examine diversification of bill shape in cavity-nesting, frugivorous birds using μCT of museum skins. Asian and African barbets (two separate clades within the Ramphastoidea; Piciformes) exhibit high species diversity and conserved morphologies (36). The Asian barbets (Megalaimidae) have diversified primarily in wetter forested habitats in South and Southeast Asia, whereas the African barbets (Lybiidae) inhabit primarily drier woodland and savanna habitats in sub-Saharan Africa (37, 38). The combination of diverse habitats, similar ecologies (including cavity excavation) and comparable species diversity render the African and Asian clades suitable to test the influence of environmental factors driving diversification of bill shape. Both clades occupy distinct climatic spaces (36), and thus putatively experience differences in the mechanical properties of the excavation substrate and the resulting impact stresses to the bill. This, in turn, may influence adaptive pressures on bill shape. However, morphological diversity in cavity-excavating birds has received very little study. In this study, we test whether bill shape variation is correlated to the climate within the ranges of these clades, as a proxy for ecological changes that could drive bill shape diversification. We will consider alternative hypotheses such as bill shape diversity accumulating as a result of phylogenetic or allometric constraints. In our approach we aim to shed light on evolutionary patterns of individual bill modules in unique avian lineages. The most exclusive clade uniting Asian and African barbets, the infraorder Ramphastoidea, also contains the Neotropical barbets and the highly divergent toucans, in which dramatic morphological shifts have occurred in the latter group (36) associated with the loss of primary cavity-excavation (37). Ecomorphological divergence between barbets and toucans has been examined in a separate study (36), and we here focused on the paleotropical clades in order to compare two groups consisting entirely of primary cavity excavators, with similar bill-body proportions but occupying distinct climatic (and therefore ecological) regimes. Thus, we could disentangle the effects of phylogeny and climatic factors on bill shape in the context of cavity-excavation.

## MATERIALS AND METHODS

### Specimens and microCT scanning

Our dataset included 135 individual specimens of 69 Barbet species: 67 individuals of 38 species belonging to the family Lybiidae and 68 individuals of 31 species belonging to the family Megalaimidae (Supplementary Data), following eBird/Clements nomenclature accessed from Birds of the World (https://birdsoftheworld.org/bow/home). Within the Lybiidae, we scanned all species except for *Gymnobucco sladeni, Lybius rubrifacies* and *Lybius chaplini*, and within the Megalaimidae, we scanned all species except for *Psilopogon eximius, Psilopogon auricularis* and *Psilopogon chersonesus* (the latter two recently split, and morphologically very similar to other species) (39). Study skins were loaned from the Field Museum of Natural History in Chicago, and the American Museum of Natural History in New York. We used a Quantum GX2 micro CT scanner (PerkinElmer Inc., Waltham, Massachusetts, USA) at an operating voltage of 90 kV and current of 88 μA, with the voxel size set at 90 μm.

### Segmentation

We imported three-dimensional scans as DICOM image stacks into Amira 6 (ThermoFisher Scientific, Waltham, Massachusetts, USA). For a few larger species, we obtained two scans of each specimen and merged them using scan registration in the multi-planar viewer. To examine the bony features separately from the keratinous rhamphotheca, we virtually dissected them using image segmentation. First, we used the thresholding tool to remove most of the soft tissue and keratin. Next, we used the paintbrush tool to manually remove any segmentation artifacts and add in any features that were missed by thresholding.

### Landmark-based geometric morphometrics

Following segmentation, we imported both the original scan (containing the rhamphotheca), and the segmented bone into Stratovan Checkpoint (Stratovan Corporation, Davis, California, USA). Because study skins frequently have damaged skulls, we focused our landmarking to only the bill. For each specimen, we placed landmarks at homologous points on the bill-at the base of the culmen, the proximal and distal ends of the nares, and the base of the tomium. Next, we placed 26 equally spaced semi-landmarks (measurement error: ±0.1 mm); 13 each on the maxilla and mandible, describing the shape of the culmen, tomium and gonys. These semi-landmarks were placed by drawing a straight line through the distal end of the nare and perpendicular to the tomium for the first and last point, followed by placing subsequent points equidistant from each other to the tip of the bill (Figure 1B). Using a similar configuration of landmarks, we digitized the shapes of both the keratinous and bony structures for each specimen. All landmarks and semi-landmarks were placed on the left side of the bill, with the exception of 17 specimens that exhibited some damage on the left side. For these specimens, the landmarks were placed on the right side, and we reflected the landmark coordinates along the mid-sagittal plane so that we could directly compare them to the others.

**Figure 1.**
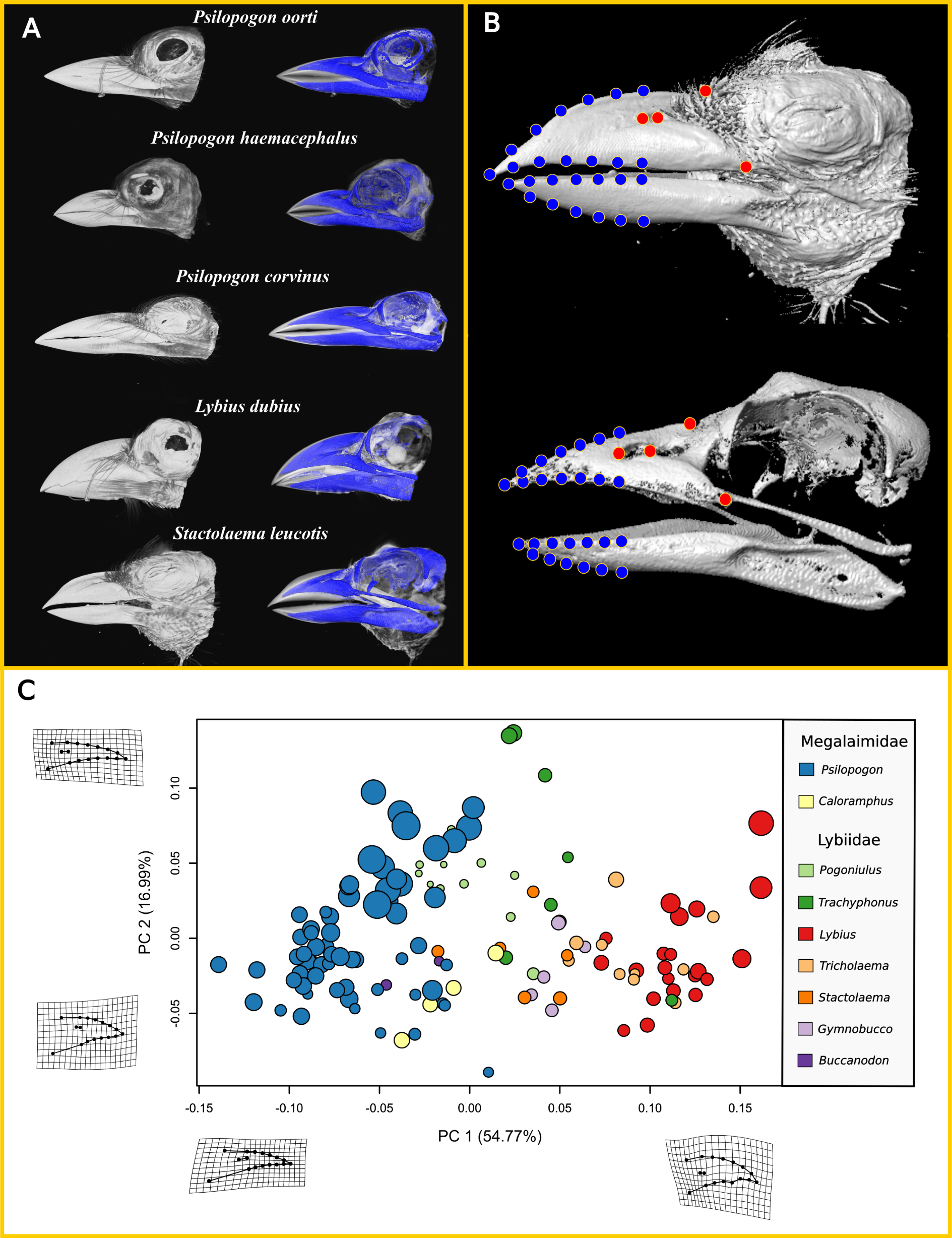
**A**. Micro-computed tomographic (CT) scans of bills from five exemplars, showing variation of the outer keratinous structures (white) and inner bony features (blue) in barbets. **B**. Segmented micro-CT scan of *Stactolaema leucotis* with landmarks (red) and semi-landmarks (blue) used for morphometric analyses. **C**. Shape morphospace for the maxillary rhamphotheca of all measured specimens, derived using non-phylogenetic principal components analysis. Each point represents a single specimen and point size and color represent centroid size and genus respectively. Asian and African barbets primarily separate along the first principal component (PC 1), which represents changes in depth and culmen curvature.

We performed all subsequent analyses in RStudio v.1.3.959 (40). To obtain shape data from the three-dimensional landmark coordinates, we conducted a Generalized Procrustes Analysis for each specimen module using the function ‘gpagen’ (in the package geomorph v3.3.1) (41). Because all landmarks were placed at or with reference to distinct anatomical loci, all of them were designated as fixed landmarks in the Procrustes transformation.

### Phylogenetic comparative analyses

We aimed to build a supermatrix by mining Genbank for as many homologous genes as possible from barbets across continental regions. Most published phylogenies have focused on the separate families and only a few had taxa from across continental regions (39, 42), some without available data (39). We used PhylotaR (43) to identify and gather all available sequences. In some cases, data from a few older studies were not picked up by PhylotaR, perhaps because of inconsistent metadata or updating of Genbank taxonomy. These were manually searched and included in our building of the supermatrix. We selected all genes that had data from at least two species from each family. If there were multiple sequences of a single species, we randomly picked one representative of each to include in our dataset. We used Muscle (44) to align individual genes and then Mesquite (45) to check alignments and concatenate the dataset. Once the supermatrix was assembled, we performed standard quality checks by comparing genetic distances and visually examining phylogenetic analyses for unusually long branches or unexpected relationships.

Our final aligned matrix included 75 taxa (72 species of Ramphastoidea and 3 outgroups) and 7563 bp from nine genes: cytochrome b (CYTB), adenosine triphosphate (ATP), fibrinogen intron 5 (FIB5), fibrinogen intron 7 (FIB7), lactate dehydrogenase (LDH), myoglobin intron 3 (MB3), muscle-specific tyrosine kinase receptor intron 4 (MUSK), NADH dehydrogenase 2 (ND2), transforming growth factor beta-1 intron 5 (TGF) bp. Supplementary Table 1 shows the complete list of species and gene information.

We used IQtree2 (46) to estimate the best partition scheme (across codon positions and genes) and model using the setting MPF+MERGE, which uses ModelFinder Plus and merges partitions that do not differ significantly in model specifications (47, 48). We estimated the phylogeny using the simultaneous tree inference and ultrafast bootstrap setting for 1000 replicates (49). To estimate a time-calibrated tree, we ran BEAST2 (50) on the same dataset. We used calibrations from Prum et al. (2015) (51) to set the ages of the root or Pici (defined by the last common ancestor of *Indicator* and *Megalaima*, now subsumed in *Psilopogon*; node 72 in Prum et al. 2015) at a mean age of 34.55 Ma (standard deviation of 12.4) and Ramphastoidea (defined by the last common ancestor of Asian, African, and Neotropical barbets; node 73 in Prum et al. 2015) at a mean age of 24.76 Ma (standard deviation of 10.2). In BEAST2, we set the site model to the same partitions as found in IQTREE with each using a HKY model of estimated gamma and substitution models. We linked clock model and tree topology across the dataset, and used a Random Local Clock model to estimate a time tree using the defined calibrations above. Next, we ran the MCMC chain for 20 million and stored every 1000 trees. We discarded the first 25% as burnin and combined the remaining runs using TreeAnnotator in the BEAST2 package. Finally, we pruned the phylogenetic tree to include only the species that were represented in the morphological dataset (Supplementary Figure 1).

Because our phylogenetic tree did not include all our sampled species, we first constructed a principal components morphospace for all specimens using the function ‘prcomp’ (stats v4.0.2) (40) to visualize overall shape variation and compare patterns observed to those in a phylomorphospace.

We used the phylogeny and Procrustes coordinates of each module to fit four models of multivariate trait evolution-Brownian motion (BM), Ornstein-Uhlenbeck (OU), early burst (EB) and lambda-through the function ‘fit_t_pl’ (RPANDA v1.8) (52). This function uses a penalized likelihood approach to fit models and a restricted maximum likelihood approach (REML) to estimate model parameters (53). We compared the model fits by extracting their respective Generalized Information Criterion (GIC) scores, using the function ‘GIC’ (mvMORPH v1.1.3) (54). Pagel’s lambda model was found to be the best fit for all four modules, and was used to inform the ensuing comparative analyses (55).

Next, we conducted a phylogenetic principal components analysis (pPCA) on the Procrustes coordinates of bill shape data using the function ‘phyl.pca_pl’ (RPANDA v1.8) (52). From these principal components, we constructed phylomorphospaces for all four modules using the function ‘phylomorphospace’ (phytools v0.7.47) (56). To statistically examine separation of the two major clades in morphospace, we performed an NPMANOVA (Non Parametric Multivariate Analysis of Variance) for each module, implemented by the function ‘adonis’ (vegan v2.5.6) (57) on all phylogenetically-corrected principal components. NPMANOVA is a non-parametric comparison of between-group dissimilarities to within-group dissimilarities (equivalent to the parametric MANOVA) based on a distance metric, using a pseudo-F ratio as a test statistic.

After examining patterns of morphospace separation, we evaluated whether this variation evolved by the integrated diversification of both the maxilla and mandible, as well as of the keratinous and bony structures. For this, we conducted pairwise phylogenetic two-block partial least squares (PPLS) regressions between the phylogenetic principal components of the four modules, using the function ‘two.b.pls’ (geomorph v3.3.1). We compared the integration levels across module pairs using the function ‘compare.pls’ (geomorph v3.3.1), which uses the z-score from a randomized distribution of r-PLS values as the effect size (41). We calculated the 95% confidence intervals for the effect size using the percentile bootstrap method.

To identify potential correlates of morphological diversification (and therefore draw inferences about its possible drivers), we carried out generalized least squares (GLS) regressions between centroid size, climate variables and bill shape (phylogenetically corrected). Climate data were obtained for each species’ range using IUCN range data (58) and the WorldClim dataset (59). We chose the mean, maximum and minimum annual temperature and precipitation as descriptors of the climatic niche, taking an average value of each parameter for each species. Temperature and precipitation represent the physical environment in which excavation substrates are formed (60), and are thus likely to influence the properties of this substrate and the biomechanical demands experienced by primary cavity-excavators. By measuring correlation of bill shape with climatic variables, we sought to obtain a preliminary understanding of the relationship between habitat differences (as discussed above) and differences in bill shape. To eliminate any multicollinearity among the chosen climatic variables, we conducted a principal components analysis on the covariance matrix of climate data using the function ‘prcomp’ (stats v4.0.2) (40). For GLS regressions, we assigned the first two principal components representing climatic conditions (PC1 and PC2) (Supplementary Data) as our explanatory variables, along with module centroid size to account for allometric variation. As our response variables, we selected the first two phylogenetically-corrected principal components (PC1 and PC2) representing module shape. All GLS regressions were performed using the function ‘gls’ (nlme v3.1) (61). In order to check for autocorrelation in our regression models, we calculated the value and significance of Blomberg’s K (a measure of phylogenetic signal) for the residuals of each regression model using the function ‘phylosig’ (phytools v0.7.47) (56).

Additionally, by mapping the PC scores onto a phylogenetic tree, we generated disparity-through-time (DTT) plots to visualize changes in within-subclade relative to between-subclade morphological disparity across time for each module. Using this, we aimed to obtain preliminary indications about whether there were differential evolutionary rate shifts across bill modules. We calculated phenotypic disparities as the average squared Euclidean distances between data points. Following this, we tested the observed DTT trajectories against a null model constructed using 1000 simulations following Brownian trait evolution on the same phylogeny. The Morphological Disparity Index (MDI), which is the area between the observed DTT curve and the median curve of the simulations, was used as a test statistic to identify significant deviations from Brownian trait evolution. We conducted the DTT analyses using the function ‘dtt’ (geiger 2.0.7) (62).

## RESULTS

### Asian and African barbets occupy distinct regions of maxillary morphospace

Micro-computed tomography enabled three-dimensional reconstruction of both bony and keratinous structures in the bills of African and Asian barbets. The keratinous rhamphotheca in both clades is expanded both rostrally and dorsoventrally past the end of the bony portion of the bill (Figure 1A). Thus, when bills are closed, the keratinous portions are in contact, and the underlying bone is away from the tip and the cutting edges of the rhamphotheca. For species possessing tomial serrations or ridged culmens (Figure 1A), these structures were not reflected in the underlying bony structures, thus there were often substantial differences in shape between these modules. The shape morphospace of the maxilla revealed that Asian and African clades generally occupy different regions of bill shape space (Figure 1C). Clades separate along the PC1 axis, which explains about 55% of total bill shape variation (Supplementary Data). The Asian barbets tend to cluster in a region of morphospace defined by a straighter, less convex culmen, whereas the African barbets generally possess a more convex culmen. The zone in morphospace where the two meet is characterized by Asian barbets with more convex maxillae (such as *Caloramphus*), and African barbets with straighter bills (for example, *Buccanodon* and *Pogoniulus*). The African genera *Lybius* and *Tricholaema* occupy a region of morphospace characterized by higher culmen curvature than other taxa.

### Barbet bills overlap in mandibular morphospace

Phylomorphospaces describing shape variation in all four modules (rhamphotheca and bone for maxilla and mandible) for the species represented in our phylogeny are shown in Figure 2. These morphospaces do not differ from the conventional morphospace shown in Figure 1C (where we represented all species that we scanned), and are thus suitable for interpretation of broad evolutionary patterns. Morphospace analyses reveal divergent patterns of morphological diversification across modules in Asian and African barbets. For both the rhamphotheca and bone, depth and culmen convexity are the primary sources of shape variation in the maxillae, as represented by the wireframes in Figure 2. The African barbets (Lybiidae) typically have deeper maxillae with arched culmens, whereas the Asian barbets (Megalaimidae) have narrower, straighter maxillae, consistent with the earlier observation. Both clades occupy distinct regions of maxillary morphospace (NPMANOVA, p<0.001, n=45) with a slightly higher degree of separation between clades for the rhamphotheca (pseudo-F = 18.25) compared to the bone (pseudo-F = 15.77). However, the two clades are not distinctly separated in the lower mandibular morphospace, for both the rhamphotheca (pseudo-F = 1.48, p = 0.196) and the bone (pseudo-F = 2.38, p = 0.06). This suggests that the diversification of bill shapes in these clades is driven primarily by changes in the shape of the maxilla, whereas the bony and keratinous structures of the mandible exhibit relatively little shape variation and the different species overlap in morphospace.

**Figure 2.**
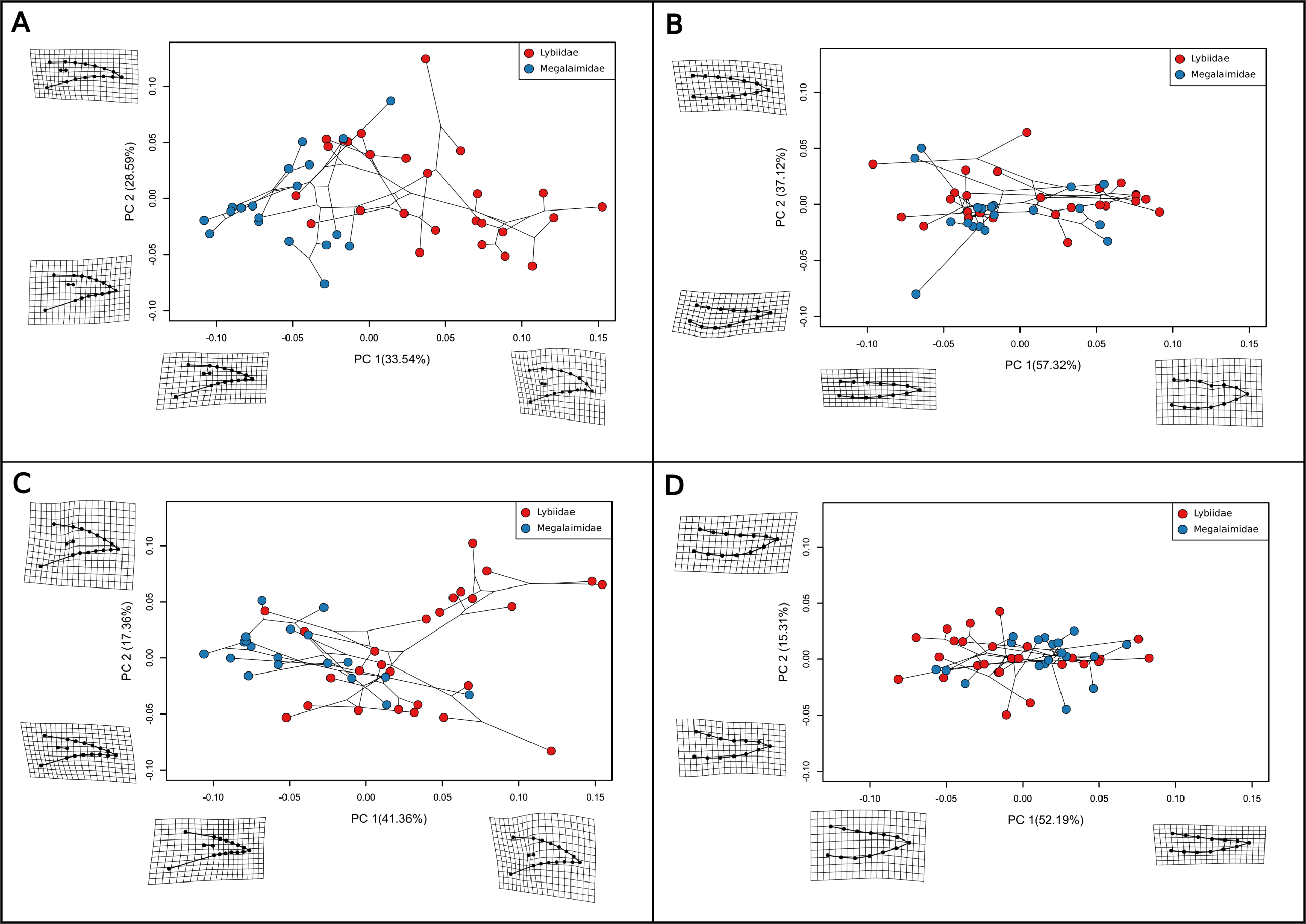
Phylomorphospaces for four bill modules- **A**. Maxillary rhamphotheca, **B**. Mandibular rhamphotheca, **C**. Maxillary bone, **D**. Mandibular bone. Phylomorphospaces depict shape variation (after controlling for phylogenetic non-independence between species in a phylogenetic principal components analysis), where each point represents a single species (blue- Asian barbets, red- African barbets). The phylogeny has been mapped on to the morphospaces, such that the connecting lines between points depict evolutionary relationships. For both the rhamphotheca and bone modules, the Asian and African barbets have distinct maxillary shapes but similar mandibular shapes. The TPS grids on the sides represent warping of bill shape along each axis with respect to the mean bill shape.

### Phenotypic integration in the diversification of barbet bills

All four modules significantly co-varied with each other (Table 1, pairwise PLS, p<0.001, n = 45) in a two-block partial least-squares analysis after correcting for phylogenetic non-independence, indicating that they are phenotypically integrated (Figure 3A-D). The highest level of integration (PLS-Z) was observed for the two maxillary modules (rhamphotheca and bone), whereas PLS-Z values for the other pairwise comparisons were broadly similar (Figure 3E). This suggests that all the four bill modules exhibit signatures of covariation in morphospace, even though the maxillary modules exhibit greater diversification in shape. The mandible rhamphotheca and bone have diversified in integration with the maxilla, albeit with less overall shape variation.

**Table 1.** Results of pairwise partial least squares (PLS) regressions between module shapes. All four bill modules strongly covary with each other.

|  | Mandibular ramphotheca | Maxillary bone |
| --- | --- | --- |
| Maxillary ramphotheca | r-PLS = 0.78, p = 0.001 | r-PLS = 0.93, p = 0.001 |
| Mandibular bone | r-PLS = 0.73, p = 0.001 | r-PLS = 0.72, p = 0.001 |

**Figure 3.**
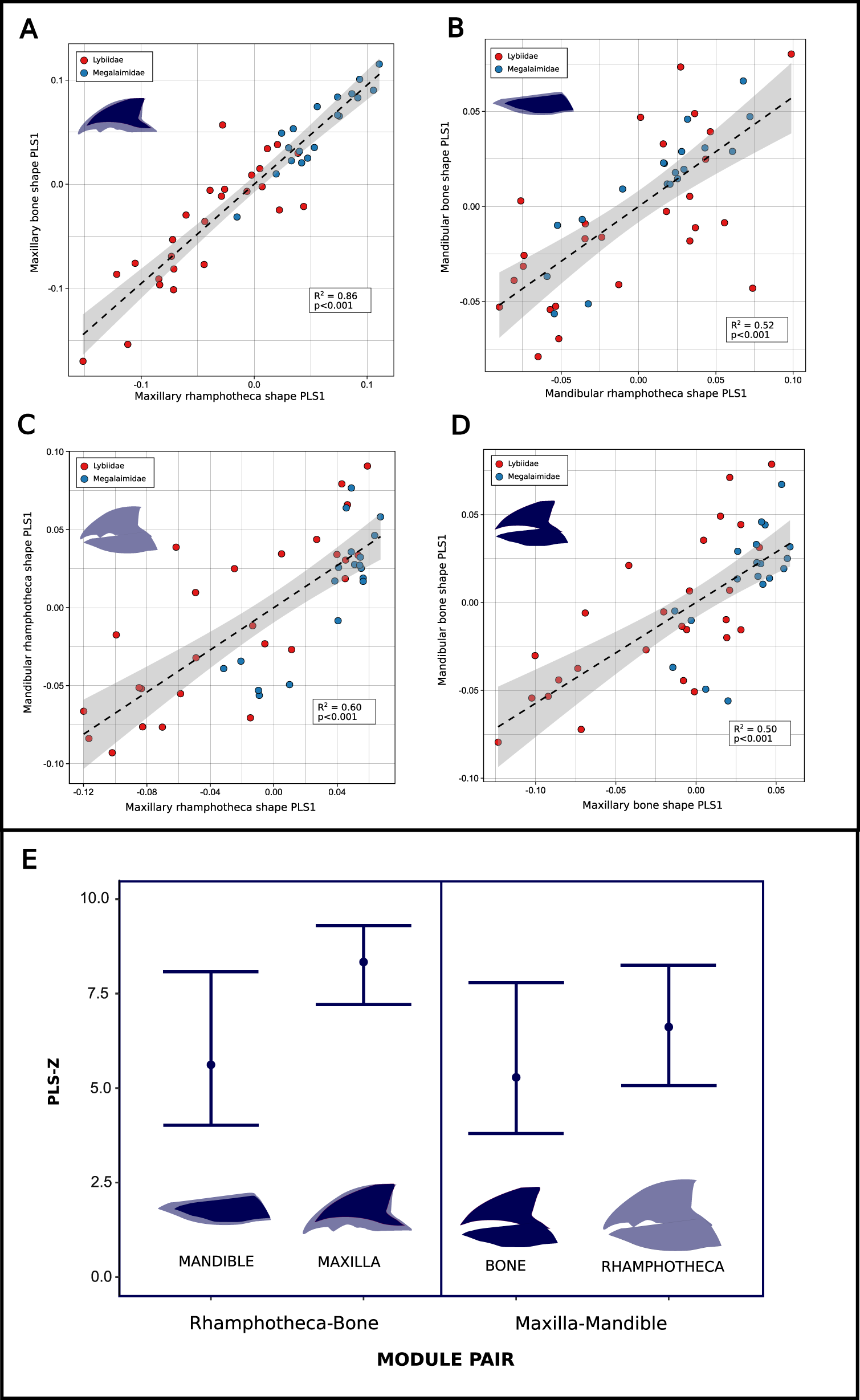
Pairwise linear regressions between module shapes, based on partial least squares (PLS) analyses on the phylogenetically corrected principal components of bill shape. **A**. Maxillary rhamphotheca and bone, **B**. Mandibular rhamphotheca and bone, **C**. Maxillary and mandibular rhamphotheca, **D**. Maxillary and mandibular bone. Each point represents a barbet species (blue- Asian, red- African). All four modules covary significantly with each other and are phenotypically integrated. **E**. Levels of phenotypic integration (depicted by the PLS-Z score) across module pairs. Confidence intervals for the PLS-Z score were generated using percentile bootstrapping.

### Putative drivers of bill shape variation

Generalized least-squares (GLS) regressions (Table 2) revealed that the first phylogenetic principal component scores representing bill shape variation in the maxillary rhamphotheca and bone were strongly predicted by climate, represented by the first two principal components PC1 (rhamphotheca: p<0.001, bone: p<0.001) and PC2 (rhamphotheca: p<0.001, bone: p<0.001) of the climate space. Both these principal components were loaded by temperature and precipitation variables, and collectively explained 74.7% of the total variation in climate. Bill centroid size was also significantly correlated with maxillary bone shape (p < 0.001) and rhamphotheca shape (p = 0.0028) indicating that allometry also influences shape variation. The second principal component for both maxillary modules was not significantly correlated with climatic variables or centroid size (p>0.1), and the mandibular modules exhibited no overall influence of climate and centroid size on shape (p>0.05 for all principal components, except between PC2 of mandibular rhamphotheca and PC2 of climate). Blomberg’s K was not statistically significant for the residuals of regressions on PC1 for both maxillary modules (rhamphotheca: K = 0.07, p = 0.65; bone: K = 0.06, p = 0.69), indicating that the observed correlations were not influenced by phylogenetic signal. Therefore, our results suggest that both ecological factors and allometric scaling exert influences on bill shape (particularly the maxilla), with the African barbets (inhabiting drier habitats) exhibiting greater culmen convexity than the forest-dwelling Asian species. In addition, within clades, changes in body size have also influenced bill shape.

**Table 2.**
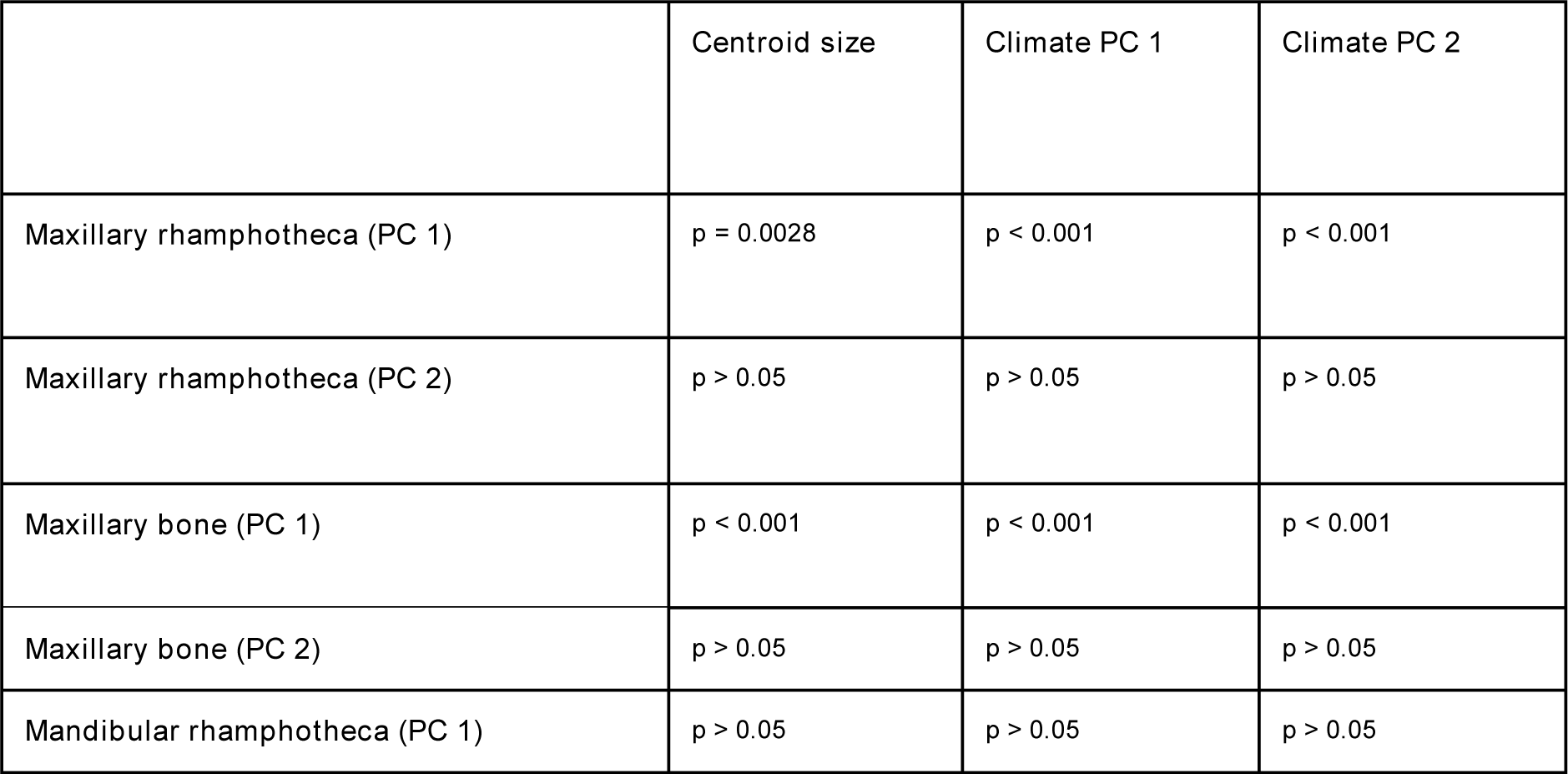
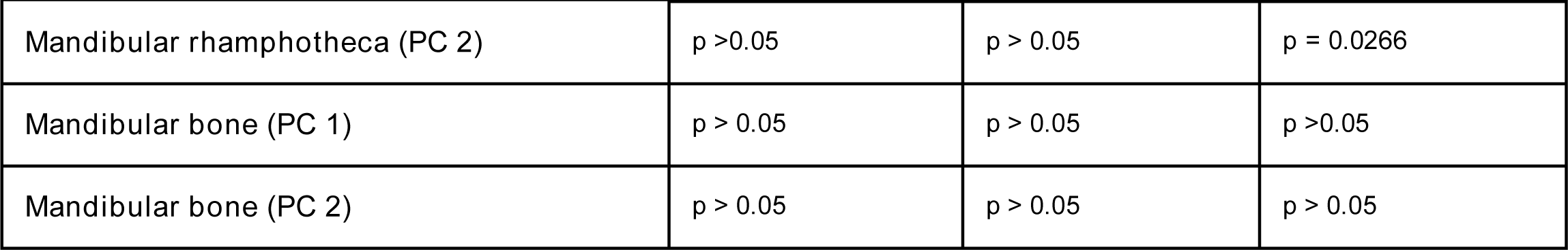
Results of generalized least squares analyses, with response variables (shape) as rows and predictor variables (centroid size and climate variables) as columns. Bold values indicate statistical significance.

|  | Centroid size | Climate PC 1 | Climate PC 2 |
| --- | --- | --- | --- |
| Maxillary ramphotheca (PC 1) | p = 0.0028 | p < 0.001 | p < 0.001 |
| Maxillary ramphotheca (PC 2) | p > 0.05 | p > 0.05 | p > 0.05 |
| Maxillary bone (PC 1) | p < 0.001 | p < 0.001 | p < 0.001 |
| Maxillary bone (PC 2) | p > 0.05 | p > 0.05 | p > 0.05 |
| Mandibular ramphotheca (PC 1) | p > 0.05 | p > 0.05 | p > 0.05 |
| **Mandibular ramphotheca (PC 2)** | p > 0.05 | p > 0.05 | **p = 0.0266** |
| **Mandibular bone (PC 1)** | p > 0.05 | p > 0.05 | p > 0.05 |
| **Mandibular bone (PC 2)** | p > 0.05 | p > 0.05 | p > 0.05 |

Disparity-through-time (DTT) plots (Figure 4) depict the relative subclade disparity through time for all four bill modules, where the solid line represents the empirical trajectory of shape evolution and the dotted line represents the mean trajectory expected under Brownian trait evolution. The empirical trajectories for all four modules remained relatively stable through time and exhibited non-significant Morphological Disparity Index (MDI) values (p-value>0.1), indicating that the pattern of morphological divergence in barbet bills does not broadly deviate from Brownian motion. For earlier analyses, we observed that a phylogenetic model incorporating Pagel’s lambda (a generalization of the Brownian motion model of evolution) was the best fit to patterns of trait evolution in barbets. Taken together, these lines of evidence are consistent with a gradual accumulation of bill shape diversity in these clades, and some late-stage diversification. The DTT also suggests a lack of “early-burst” diversification events in the evolution of barbet bills.

**Figure 4.**
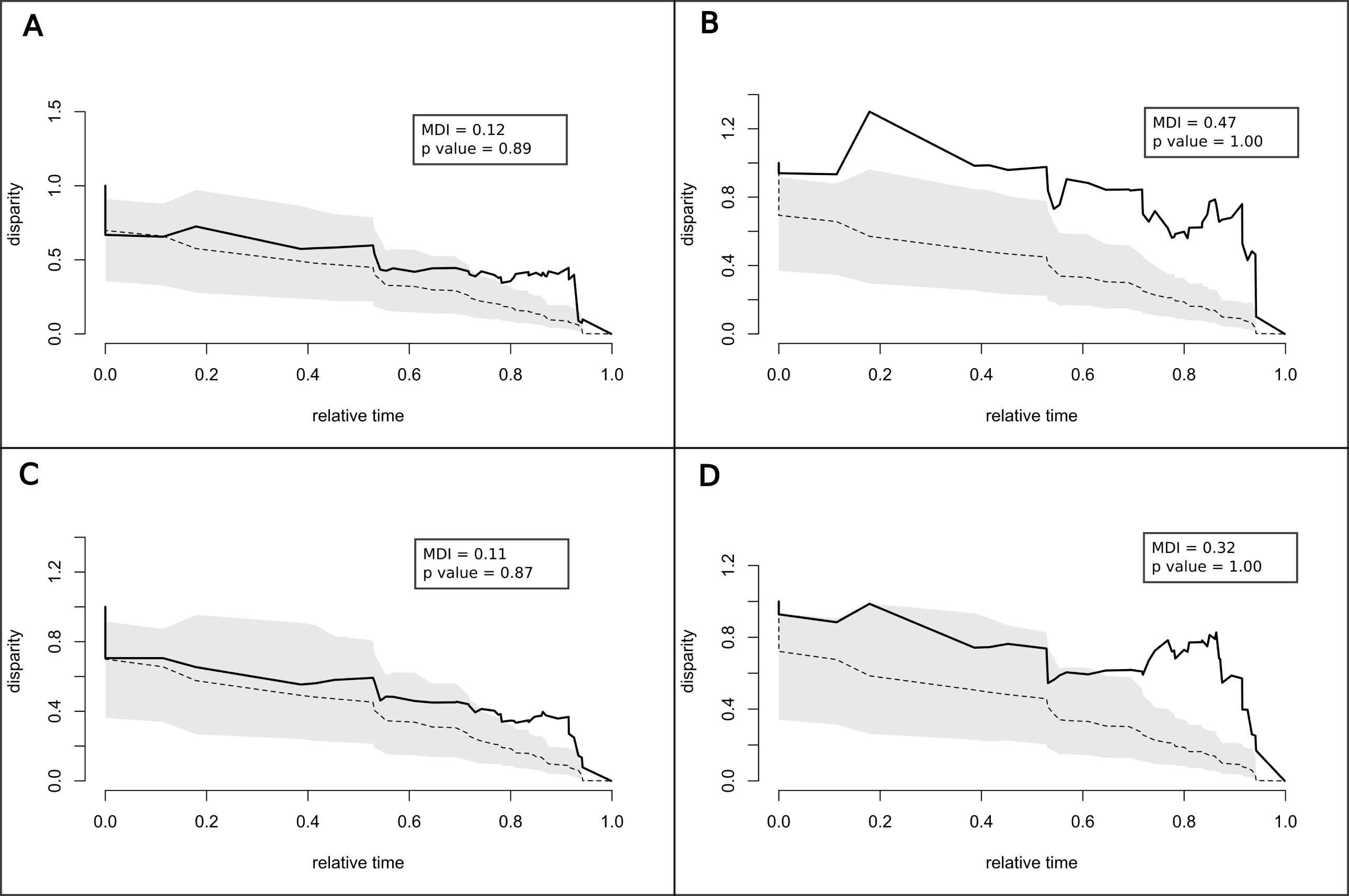
Disparity-through-time plots for all four modules **A**. Maxillary rhamphotheca, **B**. Mandibular rhamphotheca, **C**. Maxillary bone, **D**. Mandibular bone. All four modules show evidence of gradual morphological diversification without major shifts in the rate of change.

## DISCUSSION

In summary, we uncovered evidence that the bills of Asian and African barbets have diversified under distinct climatic regimes with evidence of an allometric constraint, and without evidence of an early burst of diversification. Climate variables (correlates of the ecological conditions under which these clades have evolved) are correlated with bill shape, indicating that the drier climatic regimes occupied by African barbets (37) are associated with deeper, more convex culmens. On the other hand, the wetter biotopes occupied by Asian barbets are associated with straighter, narrower culmens. Both clades have thus explored different regions of bill shape morphospace, particularly in the maxilla, whereas the mandible does not exhibit this shape divergence between continents and clades. However, we uncovered significant integration between bill modules. Thus, although different parts of the bill covary with each other, they differ in the extent of shape variation between species.

### The accumulation of bill shape diversity in Asian and African barbets

Barbets across the tropics broadly represent a conserved morphotype, with very similar body proportions across species, except in the Neotropics (63), where the toucans have diverged into a different morphological space (36). This may have resulted from ecological opportunity (64, 65), and future studies will aim to separately explore the question of ecological opportunity and its influence on bill shape and cavity nesting. The clades of Asian and African barbets, on the other hand, are relatively conserved in broad body proportions, possess roughly similar species diversity, and have evolved under broadly different climatic regimes. A large proportion of the African barbets inhabit savanna and woodland ecosystems, which are drier, whereas the Asian barbets mostly inhabit wet forest habitats (37, 38). As cavity excavators, these distinct climatic regimes are likely to exert an influence on the interactions of barbets with tree species within their habitats, as well as the availability of resources for nest or roost site excavation. Thus, the two Paleotropical barbet clades present a unique opportunity to understand how the bird bill diversifies in the absence of major morphological shifts. In spite of the differences between the maxillae of Asian and African barbets, a disparity-through-time analysis demonstrated that bill shape does not show signs of an early burst in diversification. This, combined with the conserved body proportions of most barbets, points toward a gradual accumulation of bill shape diversity in these two clades, without the major shifts seen in island adaptive radiations (6, 19). In contrast to bill shape, changes in overall body size (at least in the Asian barbets) appear to have occurred multiple times, resulting in distinct morphological classes (66).

Although we scanned nearly all species of Asian and African barbets, some of these species were missing from our phylogenetic tree. However, the phylogeny did represent more than 60% of overall barbet diversity, covering all the major subclades of Asian and African barbets (42, 67), and our phylogenetic coverage is thus comparable to that of other recent studies using CT scans (68). In addition, we note that the conventional morphospace (Figure 1C), which contained all species we scanned, did not differ from the phylogenetically corrected analyses, suggesting that the addition of further species is unlikely to alter the patterns we reported here. The six species missing from our scan dataset are members of clades and morphological groups that were well-represented in our dataset, and are also thus unlikely to alter the patterns we observed.

### Drivers and functional consequences of bill shape diversification

As mentioned above, previous studies have shown evidence of changes in the size of barbets without much change in body proportions (36). Here, we demonstrated that changes in size exert an influence on bill shape, with a significant effect of allometry after correcting for phylogenetic non-independence. In addition, we found that climatic variables representing temperature and precipitation were significantly correlated with variation in bill shape. These climatic variables are proxies of the ecological niche of an organism, and are also important drivers of ecological traits (69, 70), and this niche is expected to exert an influence on functional traits and morphological diversity. For example, because barbets are primary nest excavators (71), the availability of resources (suitable trees or logs) is likely to be influenced by the climatic conditions that drive habitat structure, the availability of dead or rotting trees for excavation, and the material properties of the excavation substrate (60), as with other cavity-nesting birds such as hornbills (72–74). Our results suggest that as a result of these ecological factors coupled to allometric scaling, Asian and African barbets have explored distinct regions of maxilla shape space. This suggests the potential for future studies to uncover the functional consequences of barbet morphology, and the role that differences in bill shape play in excavation performance, given the distinct habitat preferences of the two clades.

Barbets are largely canopy birds, feeding on soft fruit (with animal matter consumed in the breeding season) (37). The bills of barbets support the excavation of nest and roost hollows, typically in the trunk or branch of a tree. Many barbets appear to excavate in dead wood, although some also excavate in living trees (37). We hypothesize that cavity nesting has driven the evolution of the barbet bill, with variation in bill shape related to excavation performance in various habitats. However, the functional consequences of the greater convexity of African barbet bills remains unclear. These differences may result in behavioral divergence between barbet taxa, and also in the sites selected for cavity excavation as a result of the biomechanical consequences of bill shape diversity.

### Modularity and integration in the diversification of bird bills

Finally, our analyses identified significant integration across all four modules of the bill, even though the maxilla has diversified to the greatest extent. This is consistent with the bill diversifying as a single integrated unit, with different degrees of shape variation between modules. Most studies of bill shape usually only consider the shape of the outer rhamphotheca (75) or the skull module (13, 76), and there has been virtually no examination of modularity in the different structures of the bill. A recent study on kingfishers uncovered that the bone and the rhamphotheca of the bird bill evolve at different rates (68). Our study, on the other hand, uncovered similar patterns of diversification for both bone and rhamphotheca. However, by examining the maxilla and mandible as separate modules, we found that the two have diversified to different extents. By using computed tomography and virtual dissection of museum skins, which constitute the vast majority of specimen holdings in natural history collections (35), we have shown the utility of this important resource to quantify shape variation in individual bill modules.

In summary, the rhamphotheca, which generally covers and extends from the bony parts of the bill, is significantly integrated with the latter. The shapes of maxilla and the mandible are also significantly integrated with each other, albeit maxillary shape varies to a greater extent. Maxillary shape diversification appears to have been driven by ecological factors and allometric scaling, and the diversification of barbet bills has occurred without any evidence of major morphological shifts. This relatively gradual diversification has centered around changes in culmen convexity, whereas mandibular morphology has remained relatively invariant. The patterns we report here contrast with insular bird lineages, particularly adaptive radiations such as the Darwin’s finches and Hawaiian honeycreepers, where bill morphospace has diverged significantly and relatively rapidly from a common ancestor, with major morphological shifts (6, 14). It is likely that these adaptive radiations represent a release from constraints in limited geographic areas such as islands. On the other hand, on the continents of Asia and Africa, allometric constraints and integration set the stage for more subtle variation in bill shape of barbets, driven by changes in habitat and the accompanying climatic conditions. Comparing the integration and diversification of bill shape modules across diverse lineages will enable us to better understand the drivers of shape variation in this complex, multifunctional structure. In setting the stage for this process, our study demonstrates both the general significance of comparative study of natural history collections, and also the importance of studying bill shape evolution in diverse groups of birds with varied ecological and evolutionary histories.

## ACKNOWLEDGMENTS AND FUNDING

We are grateful to Nicholas Souza for scanning specimens, Thomas Sanger and James Cheverud at Loyola University Chicago for the use of the CT scanner, to Ben Marks, John Bates and Shannon Hackett at the Field Museum of Natural History, Chicago, and to Paul Sweet and Peter Capainolo at the American Museum of Natural History, New York. Finally, we thank Sanjay P. Sane and Abin Ghosh for advice on segmentation, and Chad Eliason and Helen James for useful discussions and feedback on analysis. AK was funded by an INSPIRE Faculty Award from the Department of Science and Technology, Government of India, an Early Career Research Grant (ECR/2017/001527) from the Science and Engineering Research Board (SERB), Government of India and an initiation grant from IISER Bhopal. VC is the recipient of a KVPY Fellowship from the Government of India. SR was funded by a U.S. National Science Foundation grant DEB-1457624.

