## Supplementary Table 1 for "Climate influences bill shape diversification in cavity-excavating birds"

Table 1. List of species and Genbank accession number of genes used in the supermatrix phylogeny. First column lists whether species were included in pruned tree used for comparative analyses.

| In Comp. Analyses | Species | Family | CYTB | ATP | FIB5 | FIB7 | LDH | MB3 | MUSK | ND2 | TGF |
|---|---|---|---|---|---|---|---|---|---|---|---|
| Y | *Psilopogon asiaticus* | Megalaimidae | DQ182310.1 | - | - | - | KJ455267.1 | - | - | KJ455491.1 | - |
| Y | *Psilopogon franklinii* | Megalaimidae | AY279269.1 | - | - | AY279225.1 | - | - | - | DQ479147.1 | - |
| Y | *Psilopogon haemacephalus* | Megalaimidae | AY279274.1 | - | - | AY279230.1 | KJ455268.1 | - | - | KJ455492.1 | - |
| Y | *Psilopogon chrysopogon* | Megalaimidae | MN991619.1 | - | - | - | - | - | MN991771.1 | MN991485.1 | MN991934.1 |
| | *Psilopogon haemacephalus delicus* | Megalaimidae | HQ877157.1 | - | - | HQ874501.1 | - | - | - | HQ874476.1 | - |
| | *Psilopogon haemacephalus indicus* | Megalaimidae | HQ877152.1 | - | - | HQ874499.1 | - | - | - | HQ874471.1 | - |
| | *Psilopogon haemacephalus roseus* | Megalaimidae | HQ877159.1 | - | - | HQ874502.1 | KJ455269.1 | - | - | HQ874478.1 | - |
| Y | *Psilopogon lineatus* | Megalaimidae | AY279272.1 | - | - | AY279228.1 | - | - | - | KJ455493.1 | - |
| Y | *Psilopogon malabaricus* | Megalaimidae | HQ877174.1 | - | - | HQ874517.1 | - | - | - | HQ874493.1 | - |
| Y | *Psilopogon monticola* | Megalaimidae | AY279275.1 | - | - | AY279231.1 | - | - | MN991773.1 | MN991486.1 | MN991936.1 |
| Y | *Psilopogon mystacophanos* | Megalaimidae | AF123529.1 | - | - | AY279229.1 | - | - | MN991774.1 | MN991487.1 | - |
| Y | *Psilopogon oorti* | Megalaimidae | AY279270.1 | - | - | AY279226.1 | - | - | - | - | - |
| Y | *Psilopogon faber* | Megalaimidae | DQ182314.1 | - | - | - | - | - | - | - | - |
| Y | *Psilopogon rubricapillus* | Megalaimidae | HQ877172.1 | - | - | HQ874515.1 | - | - | - | HQ874491.1 | - |
| Y | *Psilopogon virens* | Megalaimidae | AY279271.1 | - | - | AY279227.1 | KJ455270.1 | - | - | KJ455494.1 | - |
| Y | *Psilopogon viridis* | Megalaimidae | HQ877177.1 | - | - | HQ874520.1 | - | - | - | HQ874496.1 | - |
| Y | *Psilopogon zeylanicus* | Megalaimidae | KJ456336.1 | - | - | - | KJ455271.1 | - | - | KJ455495.1 | - |
| Y | *Psilopogon australis* | Megalaimidae | - | - | - | - | - | - | MN991770.1 | MN991484.1 | MN991933.1 |
| | *Psilopogon eximius* | Megalaimidae | - | - | - | - | - | - | MN991772.1 | MG546418.1 | MN991935.1 |
| Y | *Psilopogon nuchalis* | Megalaimidae | DQ182316.1 | - | - | - | - | - | - | - | - |
| Y | *Psilopogon pulcherrimus* | Megalaimidae | - | - | - | - | - | - | MN991775.1 | MN991488.1 | MN991937.1 |
| Y | *Psilopogon pyrolophus* | Megalaimidae | AF123530.1 | - | - | - | - | - | - | - | - |
| Y | *Calorhamphus fuliginosus* | Megalaimidae | AY279267.1 | - | - | AY279224.1 | - | - | MN991664.1 | MN991389.1 | MN991824.1 |
| Y | *Buccanodon duchaillui* | Lybiidae | AY279290.1 | - | - | AY279246.1 | - | - | - | - | - |
| Y | *Gymnobucco calvus* | Lybiidae | AY279276.1 | - | - | AY279232.1 | - | - | - | - | - |
| Y | *Lybius torquatus* | Lybiidae | AY279294.1 | - | - | AY279250.1 | - | - | - | - | - |
| Y | *Lybius vieilloti* | Lybiidae | AY279293.1 | - | - | AY279249.1 | - | - | - | - | - |
| Y | *Pogoniulus atroflavus* | Lybiidae | AY279283.1 | MH364256.1 | - | AY279239.1 | - | MH364317.1 | - | - | - |
| Y | *Pogoniulus bilineatus* | Lybiidae | MG437420.1 | MH364270.1 | MG576343.1 | - | - | MH364307.1 | - | - | - |
| Y | *Pogoniulus chrysoconus* | Lybiidae | AY279281.1 | MG230183.1 | MG673552.1 | AY279237.1 | - | MH364342.1 | - | - | - |
| Y | *Pogoniulus coryphaea* | Lybiidae | AY279284.1 | MH364258.1 | - | AY279240.1 | - | MH364319.1 | - | - | - |
| | *Pogoniulus leucolaimus* | Lybiidae | MH364227.1 | MG230161.1 | MG673548.1 | - | - | MH364305.1 | - | - | - |
| Y | *Pogoniulus leucomystax* | Lybiidae | MH364243.1 | MH364252.1 | - | - | - | MH364338.1 | - | - | - |
| | *Pogoniulus makawai* | Lybiidae | MG211673.1 | MG230187.1 | MG673556.1 | - | - | - | - | - | - |
| Y | *Pogoniulus pusillus* | Lybiidae | EU167002.1 | MH364277.1 | - | - | - | MH364336.1 | - | - | - |
| Y | *Pogoniulus scolopaceus* | Lybiidae | AY279282.1 | MH364255.1 | - | AY279238.1 | - | MH364318.1 | - | - | - |
| Y | *Pogoniulus simplex* | Lybiidae | MG211670.1 | MG230179.1 | MG576401.1 | - | - | MH364326.1 | - | - | - |
| Y | *Pogoniulus subsulphureus* | Lybiidae | MG211671.1 | MG230169.1 | MG576399.1 | - | - | MH364310.1 | - | - | - |
| Y | *Lybius bidentatus* | Lybiidae | AF123527.1 | - | - | - | - | - | - | - | - |
| Y | *Lybius dubius* | Lybiidae | AY279291.1 | - | - | AY279247.1 | - | - | - | - | - |
| Y | *Lybius melanopterus* | Lybiidae | AY279292.1 | - | - | AY279248.1 | - | - | - | - | - |
| Y | *Stactolaema leucotis* | Lybiidae | AY279277.1 | - | - | AY279234.1 | - | - | - | - | - |
| Y | *Stactolaema olivacea* | Lybiidae | AY279279.1 | - | - | AY279235.1 | - | AY165822.1 | - | - | - |
| Y | *Stactolaema whytii* | Lybiidae | AY279280.1 | - | - | AY279236.1 | - | - | - | - | - |
| Y | *Trachyphonus darnaudii* | Lybiidae | AF123531.1 | - | - | AY279254.1 | - | - | - | - | - |
| Y | *Trachyphonus erythrocephalus* | Lybiidae | AY279299.1 | - | - | AY279255.1 | - | - | - | - | - |
| Y | *Trachyphonus purpuratus* | Lybiidae | AY279295.1 | - | - | AY279253.1 | - | - | - | - | - |
| | *Trachyphonus usambiro* | Lybiidae | EU166982.1 | - | - | - | - | AY165825.1 | - | - | - |
| Y | *Tricholaema diademata* | Lybiidae | AY279285.1 | MG230178.1 | MG576402.1 | AY279241.1 | - | MH364325.1 | - | - | - |

| | | | | | | | | | | | |
|---|---|---|---|---|---|---|---|---|---|---|---|
| Y | *Tricholaema hirsuta* | Lybiidae | AY279286.1 | - | - | AY279243.1 | - | - | - | - | - |
| Y | *Tricholaema lachrymosa* | Lybiidae | AY279289.1 | - | - | AY279245.1 | - | - | - | - | - |
| Y | *Tricholaema leucomelas* | Lybiidae | AY279288.1 | - | - | AY279244.1 | - | - | - | - | - |
| | *Semnornis frantzii* | Semnornithidae | AF123511.1 | - | - | - | - | - | - | - | - |
| | *Semnornis ramphastinus* | Semnornithidae | AF123510.1 | - | - | AY279258.1 | - | - | - | - | - |
| | *Andigena cucullata* | Ramphastidae | AY279303.1 | - | - | AY279260.1 | - | - | - | - | - |
| | *Andigena hypoglauca* | Ramphastidae | AF123516.1 | - | - | - | - | - | - | - | - |
| | *Andigena laminirostris* | Ramphastidae | AF123515.1 | - | - | - | - | - | - | - | - |
| | *Aulacorhynchus derbianus* | Ramphastidae | AF123523.1 | - | - | - | - | - | - | - | - |
| | *Aulacorhynchus prasinus* | Ramphastidae | AY279307.1 | - | - | - | - | - | - | - | - |
| | *Pteroglossus bailloni* | Ramphastidae | AY279305.1 | - | - | - | - | - | - | - | - |
| | *Pteroglossus castanotis* | Ramphastidae | AF123520.1 | - | - | - | - | - | - | - | - |
| | *Pteroglossus inscriptus* | Ramphastidae | AY279304.1 | - | - | AY279261.1 | - | - | - | - | - |
| | *Ramphastos ariel* | Ramphastidae | AY279302.1 | - | - | AY279259.1 | - | - | - | - | - |
| | *Ramphastos culminatus* | Ramphastidae | AF123521.1 | - | - | - | - | - | - | - | - |
| | *Ramphastos cuvieri* | Ramphastidae | AF123522.1 | - | - | - | - | - | - | - | - |
| | *Selenidera gouldii* | Ramphastidae | AF123518.1 | - | - | - | - | - | - | - | - |
| | *Selenidera reinwardtii* | Ramphastidae | AY279306.1 | - | - | - | - | - | - | - | - |
| | *Selenidera spectabilis* | Ramphastidae | AF123517.1 | - | - | - | - | - | - | - | - |
| | *Capito dayi* | Capitonidae | AY279300.1 | - | - | AY279256.1 | - | - | - | - | - |
| | *Capito niger* | Capitonidae | AF123524.1 | - | - | - | - | - | - | - | - |
| | *Eubucco bourcierii* | Capitonidae | AY279301.1 | - | - | AY279257.1 | - | - | - | - | - |
| | *Eubucco richardsoni* | Capitonidae | AF123513.1 | - | - | - | - | - | - | - | - |
| | *Sphyrapicus varius* | Picidae | AF123532.1 | - | - | - | - | - | - | - | - |
| | *Chrysophlegma mentale* | Picidae | AY279265.1 | - | - | AY279221.1 | - | - | - | - | - |
| | *Indicator minor* | Indicatoridae | AY279266.1 | - | - | AY279222.1 | - | - | - | - | - |
