## Supplementary figures and images for "Climate influences bill shape diversification in cavity-excavating birds"

### Supplementary Figure 1

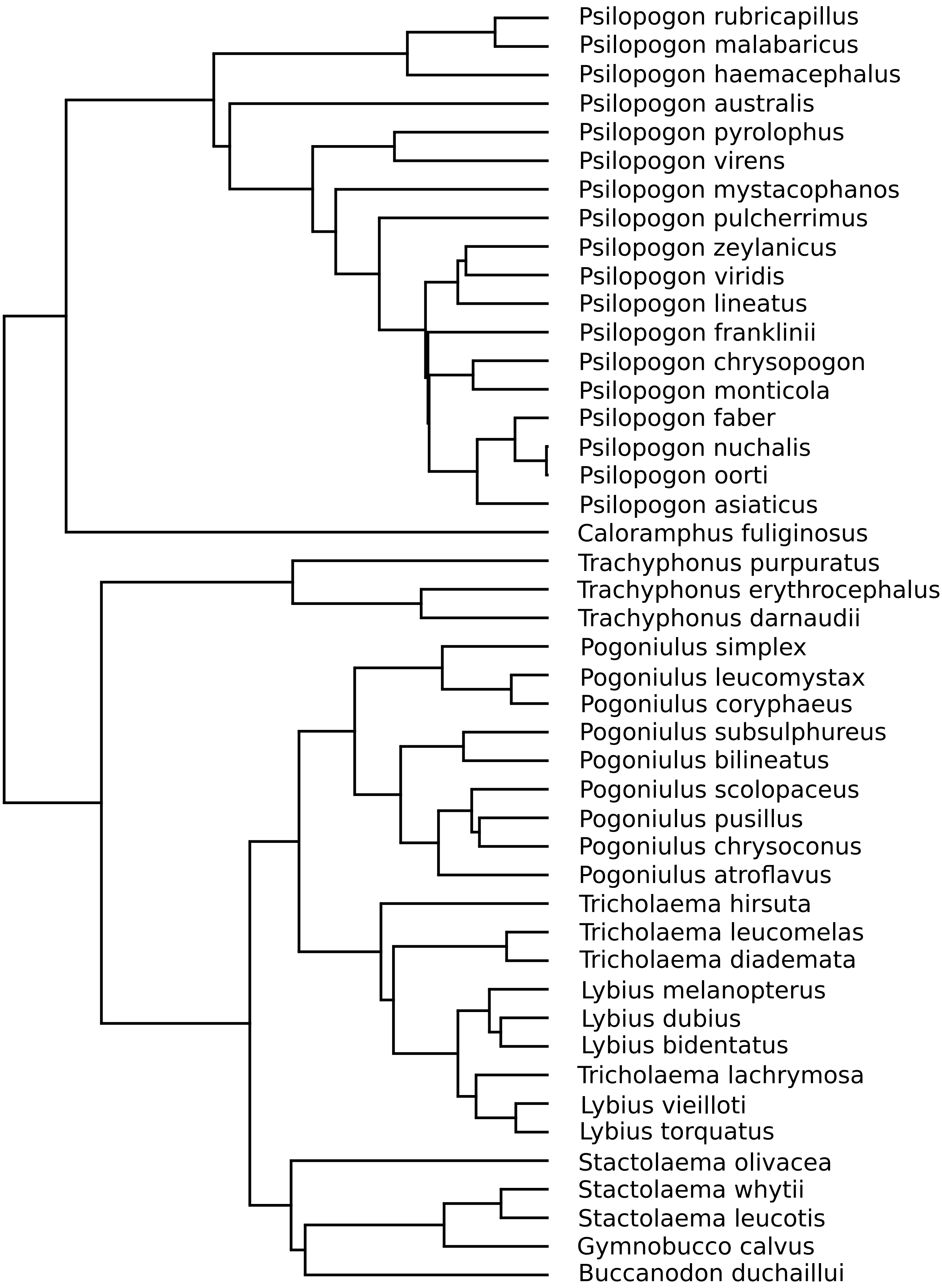
