## Supplementary Data for "Climate influences bill shape diversification in cavity-excavating birds"

**SUPPLEMENTARY DATA 1- LIST OF SPECIMENS**

**FMNH-** Field Museum of Natural History, Chicago

**AMNH-** American Museum of Natural History, New York

| **Species** | **Number of specimens** | **Museum** | **Catalogue No.** |
| --- | --- | --- | --- |
| **Lybiidae** |  |  |  |
| *Stactolaema anchietae* | 2 | FMNH | 10099 |
|  |  | FMNH | 10098 |
| *Lybius undatus* | 2 | FMNH | 82720 |
|  |  | FMNH | 82719 |
| *Lybius dubius* | 2 | FMNH | 278829 |
|  |  | FMNH | 285653 |
| *Lybius minor* | 4 | FMNH | 205249 |
|  |  | FMNH | 205250 |
|  |  | FMNH | 215548 |
|  |  | FMNH | 215549 |
| *Lybius guifsobalito* | 2 | FMNH | 113169 |
|  |  | FMNH | 113170 |
| *Lybius torquatus* | 2 | FMNH | 100097 |
|  |  | FMNH | 100096 |
| *Tricholaema melanocephala* | 2 | FMNH | 113250 |
|  |  | FMNH | 113251 |
| *Gymnobucco peli* | 2 | FMNH | 270748 |
|  |  | FMNH | 270747 |
| *Lybius melanopterus* | 2 | FMNH | 194782 |
|  |  | FMNH | 194794 |
| *Trachyphonus vaillantii* | 2 | FMNH | 100105 |
|  |  | FMNH | 100106 |
| *Lybius bidentatus* | 2 | FMNH | 113196 |
|  |  | FMNH | 113195 |
| *Stactolaema olivacea* | 2 | FMNH | 194936 |
|  |  | FMNH | 194937 |
| *Gymnobucco bonapartei* | 2 | FMNH | 113256 |
|  |  | FMNH | 113257 |
| *Tricholaema hirsuta* | 2 | FMNH | 113214 |
|  |  | FMNH | 113215 |
| *Gymnobucco calvus* | 2 | FMNH | 185349 |
|  |  | FMNH | 185348 |
| *Trachyphonus erythrocephalus* | 2 | FMNH | 113351 |
|  |  | FMNH | 113352 |
| *Tricholaema diademata* | 2 | FMNH | 103408 |
|  |  | FMNH | 103407 |
| *Tricholaema frontata* | 2 | FMNH | 205272 |
|  |  | FMNH | 205271 |
| *Pogoniulus atroflavus* | 2 | FMNH | 121853 |
|  |  | FMNH | 121854 |
| *Pogoniulus scolopaceus* | 2 | FMNH | 113347 |
|  |  | FMNH | 113346 |
| *Lybius vieilloti* | 2 | FMNH | 278831 |
|  |  | FMNH | 285658 |
| *Pogoniulus coryphaeus* | 2 | FMNH | 224409 |
|  |  | FMNH | 224408 |
| *Stactolaema leucotis* | 2 | FMNH | 194917 |
|  |  | FMNH | 194918 |
| *Lybius leucocephalus* | 2 | FMNH | 113175 |
|  |  | FMNH | 113176 |
| *Trachyphonus purpuratus* | 2 | FMNH | 113363 |
|  |  | FMNH | 113364 |
| *Pogoniulus chrysoconus* | 2 | FMNH | 113311 |
|  |  | FMNH | 113312 |
| *Buccanodon duchaillui* | 2 | FMNH | 113284 |
|  |  | FMNH | 113285 |
| *Pogoniulus subsulphureus* | 1 | FMNH | 121845 |
| *Tricholaema leucomelas* | 1 | FMNH | 214390 |
| *Lybius rolleti* | 1 | FMNH | 103404 |
| *Trachyphonus darnaudii* | 1 | FMNH | 103409 |
| *Pogoniulus simplex* | 1 | FMNH | 194986/194987 |
| *Pogoniulus leucomystax* | 1 | FMNH | 194989 |
| *Pogoniulus pusillus* | 1 | FMNH | 195048/195049 |
| *Tricholaema lachrymosa* | 1 | FMNH | 113244 |
| *Stactolaema whytii* | 1 | FMNH | 205264 |
| *Trachyphonus margaritatus* | 1 | FMNH | 121855 |
| *Pogoniulus bilineatus* | 1 | FMNH | 113313 |
| ***Megalaimidae*** |  |  |  |
| *Psilopogon oorti* | 2 | AMNH | 646978 |
|  |  | AMNH | 646976 |
| *Psilopogon annamensis* | 2 | FMNH | 90139 |
|  |  | FMNH | 90140 |
| *Psilopogon australis* | 2 | AMNH | 647059 |
|  |  | AMNH | 647060 |
| *Psilopogon duvaucelii* | 4 | FMNH | 211806 |
|  |  | FMNH | 211807 |
|  |  | FMNH | 231495 |
|  |  | FMNH | 231496 |
| *Psilopogon javensis* | 2 | FMNH | 189639 |
|  |  | AMNH | 646702 |
| *Psilopogon asiaticus* | 4 | FMNH | 78089 |
|  |  | FMNH | 78088 |
|  |  | FMNH | 211063 |
|  |  | FMNH | 221750 |
| *Caloramphus fuliginosus* | 2 | FMNH | 211801 |
|  |  | FMNH | 211799 |
| *Caloramphus hayii* | 2 | FMNH | 303070 |
|  |  | FMNH | 303072 |
| *Psilopogon zeylanicus* | 2 | FMNH | 189737 |
|  |  | FMNH | 189738 |
| *Psilopogon corvinus* | 2 | AMNH | 646655 |
|  |  | AMNH | 258944 |
| *Psilopogon faber* | 3 | AMNH | 647072 |
|  |  | AMNH | 647075 |
|  |  | FMNH | 96827 |
| *Psilopogon haemacephalus* | 4 | FMNH | 218040 |
|  |  | FMNH | 218873 |
|  |  | FMNH | 189643 |
|  |  | FMNH | 190408 |
| *Psilopogon rubricapillus* | 2 | FMNH | 98274 |
|  |  | FMNH | 257609 |
| *Psilopogon malabaricus* | 2 | FMNH | 231504 |
|  |  | FMNH | 231503 |
| *Psilopogon armillaris* | 2 | FMNH | 189641 |
|  |  | FMNH | 189642 |
| *Psilopogon pulcherrimus* | 2 | AMNH | 646904 |
|  |  | AMNH | 646906 |
| *Psilopogon franklinii* | 4 | FMNH | 303050 |
|  |  | FMNH | 303051 |
|  |  | FMNH | 221755 |
|  |  | FMNH | 221756 |
| *Psilopogon virens* | 2 | FMNH | 211069 |
|  |  | FMNH | 211070 |
| *Psilopogon faiostrictus* | 2 | FMNH | 76308 |
|  |  | FMNH | 76309 |
| *Psilopogon lineatus* | 2 | FMNH | 219140 |
|  |  | FMNH | 211067 |
| *Psilopogon monticola* | 2 | AMNH | 647003 |
|  |  | AMNH | 647005 |
| *Psilopogon incognitus* | 2 | FMNH | 90166 |
|  |  | FMNH | 90167 |
| *Psilopogon rafflesii* | 2 | FMNH | 80924 |
|  |  | FMNH | 80923 |
| *Psilopogon lagrandieri* | 2 | FMNH | 76300 |
|  |  | FMNH | 76301 |
| *Psilopogon nuchalis* | 2 | FMNH | 56337 |
|  |  | FMNH | 56338 |
| *Psilopogon viridis* | 2 | FMNH | 231406 |
|  |  | FMNH | 231405 |
| *Psilopogon henricii* | 2 | AMNH | 646889 |
|  |  | FMNH | 98708 |
| *Psilopogon pyrolophus* | 1 | FMNH | 363878 |
| *Psilopogon chrysopogon* | 2 | FMNH | 98702 |
|  |  | FMNH | 98703 |
| *Psilopogon mystacophanos* | 1 | FMNH | 80922 |
| *Psilopogon flavifrons* | 1 | FMNH | 256781 |

**SUPPLEMENTARY DATA 2- PRINCIPAL COMPONENTS ANALYSES DATA**

2.1 PCA on Procrustes shape variables for maxillary rhamphotheca (all specimens)

|  | **Proportion of variance** | **Cumulative proportion** |
| --- | --- | --- |
| **PC 1** | 54.76945258 | 54.76945258 |
| **PC 2** | 16.98889287 | 71.75834545 |
| **PC 3** | 8.426543945 | 80.18488939 |
| **PC 4** | 5.227190774 | 85.41208016 |
| **PC 5** | 2.958251442 | 88.37033161 |
| **PC 6** | 1.977041785 | 90.34737339 |
| **PC 7** | 1.766425334 | 92.11379872 |
| **PC 8** | 1.464484351 | 93.57828308 |
| **PC 9** | 1.410618647 | 94.98890172 |
| **PC 10** | 0.932545859 | 95.92144758 |
| **PC 11** | 0.686493501 | 96.60794108 |
| **PC 12** | 0.530923272 | 97.13886436 |
| **PC 13** | 0.442873526 | 97.58173788 |
| **PC 14** | 0.336345772 | 97.91808365 |
| **PC 15** | 0.310198614 | 98.22828227 |
| **PC 16** | 0.259823919 | 98.48810619 |
| **PC 17** | 0.238361443 | 98.72646763 |
| **PC 18** | 0.200672006 | 98.92713964 |
| **PC 19** | 0.181715109 | 99.10885475 |
| **PC 20** | 0.13013843 | 99.23899318 |
| **PC 21** | 0.120854123 | 99.3598473 |
| **PC 22** | 0.100276809 | 99.46012411 |
| **PC 23** | 0.092784906 | 99.55290901 |
| **PC 24** | 0.075310104 | 99.62821912 |
| **PC 25** | 0.05531253 | 99.68353165 |
| **PC 26** | 0.048093405 | 99.73162505 |
| **PC 27** | 0.04053808 | 99.77216313 |
| **PC 28** | 0.03476595 | 99.80692908 |
| **PC 29** | 0.02839372 | 99.8353228 |
| **PC 30** | 0.022793601 | 99.8581164 |
| **PC 31** | 0.022200432 | 99.88031684 |
| **PC 32** | 0.01810433 | 99.89842117 |
| **PC 33** | 0.014828165 | 99.91324933 |
| **PC 34** | 0.013682579 | 99.92693191 |
| **PC 35** | 0.011516319 | 99.93844823 |
| **PC 36** | 0.010808349 | 99.94925658 |
| **PC 37** | 0.009419045 | 99.95867562 |
| **PC 38** | 0.008994864 | 99.96767049 |
| **PC 39** | 0.006952525 | 99.97462301 |
| **PC 40** | 0.006905729 | 99.98152874 |
| **PC 41** | 0.00562276 | 99.9871515 |
| **PC 42** | 0.005070989 | 99.99222249 |
| **PC 43** | 0.004122857 | 99.99634535 |
| **PC 44** | 0.003654654 | 100 |
| **PC 45** | 7.82E-27 | 100 |
| **PC 46** | 8.12E-28 | 100 |
| **PC 47** | 5.26E-28 | 100 |
| **PC 48** | 2.44E-28 | 100 |
| **PC 49** | 9.27E-29 | 100 |
| **PC 50** | 2.69E-29 | 100 |
| **PC 51** | 2.20E-29 | 100 |

2.2 Phylogenetic PCA for maxillary rhamphotheca

|  | **Proportion of variance** | **Cumulative proportion** |
| --- | --- | --- |
| **PC 1** | 33.54453746 | 33.54453746 |
| **PC 2** | 28.58801611 | 62.13255357 |
| **PC 3** | 11.86907163 | 74.0016252 |
| **PC 4** | 5.72226793 | 79.72389313 |
| **PC 5** | 4.613745556 | 84.33763868 |
| **PC 6** | 3.241611733 | 87.57925042 |
| **PC 7** | 2.348864599 | 89.92811502 |
| **PC 8** | 1.708279917 | 91.63639493 |
| **PC 9** | 1.61122578 | 93.24762071 |
| **PC 10** | 1.387693809 | 94.63531452 |
| **PC 11** | 0.831750111 | 95.46706463 |
| **PC 12** | 0.646321093 | 96.11338573 |
| **PC 13** | 0.518326139 | 96.63171186 |
| **PC 14** | 0.516136826 | 97.14784869 |
| **PC 15** | 0.287551758 | 97.43540045 |
| **PC 16** | 0.219437442 | 97.65483789 |
| **PC 17** | 0.208443837 | 97.86328173 |
| **PC 18** | 0.172740388 | 98.03602212 |
| **PC 19** | 0.157206935 | 98.19322905 |
| **PC 20** | 0.133562944 | 98.326792 |
| **PC 21** | 0.113135228 | 98.43992722 |
| **PC 22** | 0.103204684 | 98.54313191 |
| **PC 23** | 0.081455193 | 98.6245871 |
| **PC 24** | 0.079481286 | 98.70406839 |
| **PC 25** | 0.072355558 | 98.77642394 |
| **PC 26** | 0.06363533 | 98.84005927 |
| **PC 27** | 0.059115291 | 98.89917457 |
| **PC 28** | 0.056003709 | 98.95517827 |
| **PC 29** | 0.052176378 | 99.00735465 |
| **PC 30** | 0.051078125 | 99.05843278 |
| **PC 31** | 0.049771806 | 99.10820458 |
| **PC 32** | 0.047808656 | 99.15601324 |
| **PC 33** | 0.04725837 | 99.20327161 |
| **PC 34** | 0.046072029 | 99.24934364 |
| **PC 35** | 0.045615406 | 99.29495904 |
| **PC 36** | 0.045094463 | 99.34005351 |
| **PC 37** | 0.044883806 | 99.38493731 |
| **PC 38** | 0.044624143 | 99.42956146 |
| **PC 39** | 0.044318836 | 99.47388029 |
| **PC 40** | 0.044081572 | 99.51796186 |
| **PC 41** | 0.044045332 | 99.5620072 |
| **PC 42** | 0.043905324 | 99.60591252 |
| **PC 43** | 0.04385554 | 99.64976806 |
| **PC 44** | 0.043794601 | 99.69356266 |
| **PC 45** | 0.043776763 | 99.73733942 |
| **PC 46** | 0.043776763 | 99.78111619 |
| **PC 47** | 0.043776763 | 99.82489295 |
| **PC 48** | 0.043776763 | 99.86866971 |
| **PC 49** | 0.043776763 | 99.91244647 |
| **PC 50** | 0.043776763 | 99.95622324 |
| **PC 51** | 0.043776763 | 100 |

2.3 Phylogenetic PCA for maxillary bone

|  | **Proportion of variance** | **Cumulative proportion** |
| --- | --- | --- |
| **PC 1** | 41.36022997 | 41.36022997 |
| **PC 2** | 17.35655135 | 58.71678132 |
| **PC 3** | 11.84630373 | 70.56308505 |
| **PC 4** | 6.51783297 | 77.08091802 |
| **PC 5** | 4.804430313 | 81.88534833 |
| **PC 6** | 3.911598583 | 85.79694691 |
| **PC 7** | 2.780856463 | 88.57780338 |
| **PC 8** | 2.254245524 | 90.8320489 |
| **PC 9** | 1.656562355 | 92.48861126 |
| **PC 10** | 1.234462753 | 93.72307401 |
| **PC 11** | 0.939701356 | 94.66277536 |
| **PC 12** | 0.773355314 | 95.43613068 |
| **PC 13** | 0.663256216 | 96.09938689 |
| **PC 14** | 0.509744615 | 96.60913151 |
| **PC 15** | 0.361494331 | 96.97062584 |
| **PC 16** | 0.30323711 | 97.27386295 |
| **PC 17** | 0.215940204 | 97.48980315 |
| **PC 18** | 0.184520413 | 97.67432357 |
| **PC 19** | 0.145737819 | 97.82006139 |
| **PC 20** | 0.135411539 | 97.95547293 |
| **PC 21** | 0.121008473 | 98.0764814 |
| **PC 22** | 0.109794504 | 98.1862759 |
| **PC 23** | 0.096546708 | 98.28282261 |
| **PC 24** | 0.07942448 | 98.36224709 |
| **PC 25** | 0.078764726 | 98.44101182 |
| **PC 26** | 0.074646 | 98.51565782 |
| **PC 27** | 0.072208492 | 98.58786631 |
| **PC 28** | 0.067646783 | 98.65551309 |
| **PC 29** | 0.067239202 | 98.72275229 |
| **PC 30** | 0.064207892 | 98.78696019 |
| **PC 31** | 0.063410502 | 98.85037069 |
| **PC 32** | 0.061463933 | 98.91183462 |
| **PC 33** | 0.060787793 | 98.97262241 |
| **PC 34** | 0.059512769 | 99.03213518 |
| **PC 35** | 0.058887765 | 99.09102295 |
| **PC 36** | 0.058675052 | 99.149698 |
| **PC 37** | 0.057983357 | 99.20768136 |
| **PC 38** | 0.057596218 | 99.26527757 |
| **PC 39** | 0.05723838 | 99.32251595 |
| **PC 40** | 0.05698063 | 99.37949658 |
| **PC 41** | 0.056585097 | 99.43608168 |
| **PC 42** | 0.056473436 | 99.49255512 |
| **PC 43** | 0.056398699 | 99.54895382 |
| **PC 44** | 0.056382848 | 99.60533666 |
| **PC 45** | 0.056380477 | 99.66171714 |
| **PC 46** | 0.056380477 | 99.71809762 |
| **PC 47** | 0.056380477 | 99.77447809 |
| **PC 48** | 0.056380477 | 99.83085857 |
| **PC 49** | 0.056380477 | 99.88723905 |
| **PC 50** | 0.056380477 | 99.94361952 |
| **PC 51** | 0.056380477 | 100 |

2.4 Phylogenetic PCA for mandibular rhamphotheca

|  | **Proportion of variance** | **Cumulative proportion** |
| --- | --- | --- |
| **PC 1** | 57.31746 | 57.31746 |
| **PC 2** | 13.11979 | 70.43726 |
| **PC 3** | 11.2497 | 81.68696 |
| **PC 4** | 4.455829 | 86.14278 |
| **PC 5** | 3.135464 | 89.27825 |
| **PC 6** | 2.073591 | 91.35184 |
| **PC 7** | 1.788601 | 93.14044 |
| **PC 8** | 1.113655 | 94.2541 |
| **PC 9** | 0.973636 | 95.22773 |
| **PC 10** | 0.809863 | 96.0376 |
| **PC 11** | 0.557474 | 96.59507 |
| **PC 12** | 0.394705 | 96.98977 |
| **PC 13** | 0.303306 | 97.29308 |
| **PC 14** | 0.274095 | 97.56718 |
| **PC 15** | 0.218228 | 97.7854 |
| **PC 16** | 0.183971 | 97.96937 |
| **PC 17** | 0.150582 | 98.11996 |
| **PC 18** | 0.134806 | 98.25476 |
| **PC 19** | 0.123316 | 98.37808 |
| **PC 20** | 0.118398 | 98.49648 |
| **PC 21** | 0.100856 | 98.59733 |
| **PC 22** | 0.098669 | 98.696 |
| **PC 23** | 0.092929 | 98.78893 |
| **PC 24** | 0.090208 | 98.87914 |
| **PC 25** | 0.084377 | 98.96352 |
| **PC 26** | 0.082805 | 99.04632 |
| **PC 27** | 0.078085 | 99.12441 |
| **PC 28** | 0.0773 | 99.20171 |
| **PC 29** | 0.075514 | 99.27722 |
| **PC 30** | 0.074288 | 99.35151 |
| **PC 31** | 0.073524 | 99.42503 |
| **PC 32** | 0.072915 | 99.49795 |
| **PC 33** | 0.071722 | 99.56967 |
| **PC 34** | 0.071722 | 99.64139 |
| **PC 35** | 0.071722 | 99.71311 |
| **PC 36** | 0.071722 | 99.78483 |
| **PC 37** | 0.071722 | 99.85656 |
| **PC 38** | 0.071722 | 99.92828 |
| **PC 39** | 0.071722 | 100 |

2.5 Phylogenetic PCA for mandibular bone

|  | **Proportion of variance** | **Cumulative proportion** |
| --- | --- | --- |
| **PC 1** | 52.18846 | 52.18846 |
| **PC 2** | 15.31445 | 67.50291 |
| **PC 3** | 7.487338 | 74.99025 |
| **PC 4** | 5.15625 | 80.1465 |
| **PC 5** | 3.882787 | 84.02929 |
| **PC 6** | 2.569877 | 86.59917 |
| **PC 7** | 2.072872 | 88.67204 |
| **PC 8** | 1.732871 | 90.40491 |
| **PC 9** | 1.510357 | 91.91527 |
| **PC 10** | 0.942879 | 92.85814 |
| **PC 11** | 0.636112 | 93.49426 |
| **PC 12** | 0.56299 | 94.05725 |
| **PC 13** | 0.512608 | 94.56986 |
| **PC 14** | 0.479737 | 95.04959 |
| **PC 15** | 0.397222 | 95.44682 |
| **PC 16** | 0.378759 | 95.82557 |
| **PC 17** | 0.324703 | 96.15028 |
| **PC 18** | 0.281492 | 96.43177 |
| **PC 19** | 0.263038 | 96.69481 |
| **PC 20** | 0.248179 | 96.94299 |
| **PC 21** | 0.209636 | 97.15262 |
| **PC 22** | 0.194333 | 97.34696 |
| **PC 23** | 0.191904 | 97.53886 |
| **PC 24** | 0.178243 | 97.7171 |
| **PC 25** | 0.173567 | 97.89067 |
| **PC 26** | 0.169903 | 98.06057 |
| **PC 27** | 0.16356 | 98.22413 |
| **PC 28** | 0.160708 | 98.38484 |
| **PC 29** | 0.156277 | 98.54112 |
| **PC 30** | 0.151438 | 98.69256 |
| **PC 31** | 0.149677 | 98.84223 |
| **PC 32** | 0.148404 | 98.99064 |
| **PC 33** | 0.144195 | 99.13483 |
| **PC 34** | 0.144195 | 99.27903 |
| **PC 35** | 0.144195 | 99.42322 |
| **PC 36** | 0.144195 | 99.56742 |
| **PC 37** | 0.144195 | 99.71161 |
| **PC 38** | 0.144195 | 99.85581 |
| **PC 39** | 0.144195 | 100 |

2.6 PCA on climate variables

2.6.1 Proportion of variance explained

|  | **Proportion of Variance** | **Cumulative proportion** |
| --- | --- | --- |
| **PC1** | 45.42366176 | 45.42366176 |
| **PC2** | 29.23037028 | 74.65403204 |
| **PC3** | 17.46316951 | 92.11720155 |
| **PC4** | 5.449320631 | 97.56652218 |
| **PC5** | 1.767326666 | 99.33384885 |
| **PC6** | 0.666151152 | 100 |

2.6.2 Eigenvectors

|  | **PC1** | **PC2** | **PC3** | **PC4** | **PC5** | **PC6** |
| --- | --- | --- | --- | --- | --- | --- |
| **Temp_mean** | -0.32416 | -0.4567 | 0.517991 | -0.32993 | 0.532245 | 0.160897 |
| **Temp_min** | -0.44904 | -0.4624 | -0.05968 | -0.23331 | -0.70557 | -0.16949 |
| **Temp_max** | 0.349153 | -0.3236 | 0.584288 | 0.60396 | -0.25926 | -2.62E-05 |
| **Prec_mean** | -0.50479 | 0.334363 | 0.235807 | 0.305023 | 0.144911 | -0.68099 |
| **Prec_min** | -0.55954 | 0.192119 | -0.03982 | 0.434065 | -0.07198 | 0.674412 |
| **Prec_max** | 0.058986 | 0.569374 | 0.574056 | -0.43645 | -0.35425 | 0.163742 |
